# AllerStat: Finding Statistically Significant Allergen-Specific Patterns in Protein Sequences by Machine Learning

**DOI:** 10.1101/2021.08.17.456743

**Authors:** Kento Goto, Norimasa Tamehiro, Takumi Yoshida, Hiroyuki Hanada, Takuto Sakuma, Reiko Adachi, Kazunari Kondo, Ichiro Takeuchi

## Abstract

Cutting-edge technologies such as genome editing and synthetic biology allow us to produce novel foods and functional proteins. However, their toxicity and allergenicity must be accurately evaluated. Allergic reactions are caused by specific amino-acid sequences in proteins (Allergen Specific Patterns, ASPs), of which, many remain undiscovered. In this study, we introduce a data-driven approach and a machine-learning (ML) method to find undiscovered ASPs. The proposed method enables an exhaustive search for amino-acid sub-sequences whose frequencies are statistically significantly higher in allergenic proteins. As a proof-of-concept (PoC), we created a database containing 21,154 proteins of which the presence or absence allergic reactions are already known, and the proposed method was applied to the database. The detected ASPs in the PoC study were consistent with known biological findings, and the allergenicity prediction accuracy using the detected ASPs was higher than extant approaches.

**Teaser:** We propose a computational method for finding statistically significant allergen-specific amino-acid sequences in proteins.

## Introduction

Food allergies are atopic disorders, which can be classified into immunoglobulin E (IgE)- mediated and non-IgE-mediated disorders. Allergies to cow’s milk, egg, wheat, and peanuts for examples, are IgE-mediated. Food allergies are typically caused by hypersensitivity of the immune system to specific proteins in foods, which brings about various allergic reactions ranging from itchiness, swelling of the tongue, vomiting, diarrhea, hives, trouble breathing, low blood pressure, and systematic anaphylaxis in severe cases (*1, 2*). Gupta et al. (*3*) showed that children under 18 years in the United States yielded an estimated food allergy prevalence of 8%, and that approximately 40% of patients with food allergies have experienced a life-threatening allergic reaction. Food antigen-specific IgE antibodies play a pivotal role in most of immediate hypersensitivity to food components. The cross-linking of IgE receptors on mast cells and basophils triggers the release of mediators such as histamines and proteases (*4*). Antigen-specific IgE production from B cells requires for T-cell help, and T-cell-derived cytokine induces the class-switch of B cells into IgE-producing plasma cells (*5*). The production of such cytokines from helper T cells is regulated by T-cell receptors, which mediate signaling through the affinity to T-cell epitope peptides presented on human leukocyte antigen (HLA) molecules expressed in professional antigen-presenting cells (APCs), and thymic microenvironments formed by medullary thymic epithelial cells (mTECs) enable T-cells lineage commitment by presenting self-antigens (*6*).

Some of proteins contained in foods are the cause of food allergies. Currently, the allergenicity of proteins has been evaluated by a method based on the homogeneity to known allergenic proteins. In many countries, the method according to the Food and Agriculture Organization (FAO)–World Health Organization (WHO) criterion is used, in which proteins having *>*35% similarity in the 80-amino-acid sliding window of allergen proteins or those identical to six-to-eight contiguous amino acids that are contained in allergen proteins are regarded as allergenic proteins (*7*). However, novel foods and functional proteins can now be created using genome editing technologies and synthetic biology. Because there is a possibility that newly produced proteins much less homogenous to known allergenic proteins may have allergenicity, more reliable prediction methods are needed. Recently, machine learning (ML) has been applied to various research field, and various bioinformatic methods have also been proposed to improve the prediction accuracy of FAO/WHO rules. Most of them are built on whether the protein contains a similar amino-acid subsequence to known allergen-specific peptides, which we call Allergen-Specific Patterns (ASPs) (*8–16*). Unfortunately, these bioinformatic methods are still limited in their ability to identify allergen proteins that does not contain known ASPs, therefore, we developed a machine learning method to identify unknown ASPs by data-driven approach.

The goal of this study is to identify new ASPs that are responsible for allergenicity from a database of allergen proteins and non-allergen proteins. We propose an ML method called *aller-Stat* to efficiently identify statistically reliable ASPs, and utilize the results to investigate the biological mechanisms of allergic reactions and predicting food allergic reactions. This problem is both computationally and statistically challenging. The computational challenge is that, since the number of all possible sub-sequences in protein amino-acid sequences is extremely huge, we need to develop a computational trick that enables an efficient exhaustive search. The statistical challenge is that, when a selection is made from a huge number of candidates, it is difficult to properly evaluate the reliability (*p*-values and confidence intervals) of the selection, due to *selection bias* (*c*.*f*., multiple-comparison bias). Hence, it is necessary to develop a method to properly mitigate the bias. The main novelty of allerStat is that it overcomes these computational and statistical challenges by effectively combining sequence mining (*17–24*) and multiple testing correction methods (*25–37*).

As a proof-of-concept (PoC) study, we developed a dataset consisting of 21,154 proteins (2,248 allergen and 18,906 non-allergen proteins) by collecting them from multiple databases. We applied allerStat to the dataset and successfully identified 5,994 statistically significant ASPs after correcting the selection bias at the significance level of *α* = 0.05. We observe that a part of the identified ASPs have high HLA type II binding activity which plays a significant role in atopic diseases and food allergies (*38, 39*) and are consistent with known Inger epitopes (*40, 41*). Furthermore, we develop an allergic reaction prediction method based on the identified ASPs and demonstrate that its prediction accuracy of the method is better than existing methods (*42–44*). We provide the database for the PoC study in the supplementary materials and the code is provided at https://github.com/takeuchi-lab/allerStat.

## Results

In this paper we propose a ML method called allerStat for identifying amino-acid subsequences highly associated with allergic reactions, which we call ASPs. As a PoC, we constructed a protein dataset that contains both allergic and non-allergic proteins in various biological categories, and we applies allerStat to the dataset.

### Allergen Protein Dataset

The input format of the dataset for allerStat is illustrated in Fig. 1(**A**). Each row of the table represents an individual protein. Each protein consists of three pieces of information: amino-acid sequence, biological category, and presence or absence of allergic reaction (label). We call a protein that causes or does not cause allergic reaction as an *allergenic protein* or a *non-allergenic protein*, respectively. Biological category information is needed to avoid misidentifying category-specific patterns as ASPs. From the original dataset, we extracted the information as illustrated in Fig. 1(**B**), in which each biological category in the dataset is classified into three types. The first type contains both allergen and non-allergenic proteins (called *paired category*), the second type only contains allergenic proteins (called *positive-only category*), and the third type only contains non-allergenic proteins (called *negative-only category*). The distinction of these three types of categories is needed because, e.g., amino-acid subsequences frequently observed in a positive-only category cannot be distinguished whether they are specific to the category or to an allergic reaction.

**Fig. 1.**
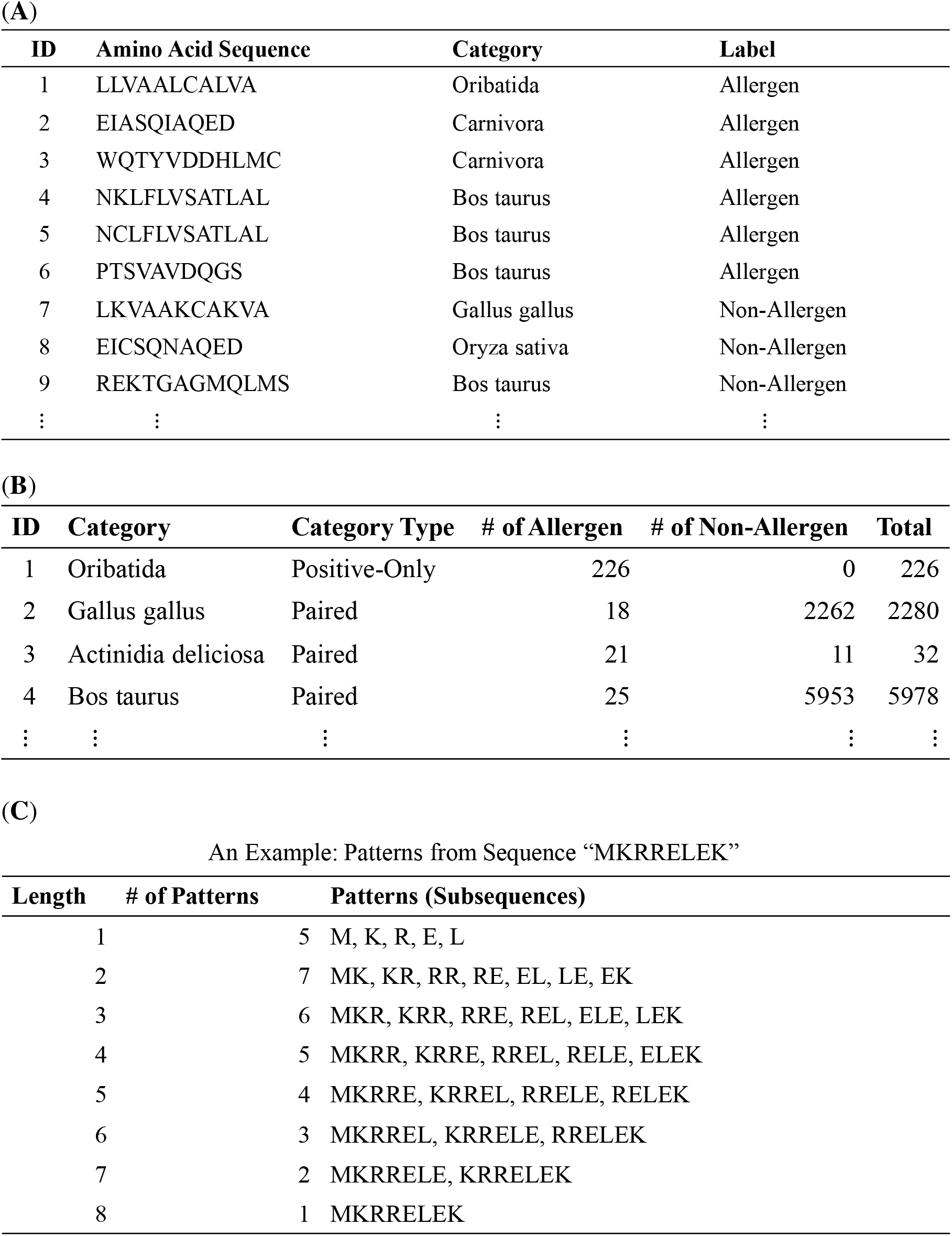
Illustration of the data format for the proposed method allerStat. (**A**) Input data format. Each line indicates a protein, and each protein is represented by an amino-acid sequence, biological category name, and allergenic reaction status. (**B**) Data format for biological categories. Three types of categories (i.e., paired category, positive-only category and negative-only category) are differently treated by the data analysis method. (**C**) Examples of patterns. We call a contiguous amino-acid subsequence a *pattern*. The figure indicates that there are 33 patterns within the amino-acid sequence MKRRELEK.

As a PoC, we developed a dataset of 21,154 proteins whose presence or absence of allergic reactions have been already verified (see the Methods section). The whole dataset in the form of Fig. 1(**A**) is given in Data S1 and the biological category information of the dataset in the form of Fig. 1(**B**) is given in Data S2. The PoC dataset consists of 2,248 allergenic proteins and 18,906 non-allergenic proteins. The average, minimum and maximum lengths of the proteins are 421, 5, and 34,350, respectively. There are 20 paired categories, 204 positive-only categories, and one negative-only category. It is generally difficult to discriminate between allergenic and non-allergenic proteins. Thus, we focused on 20 foods that have been well analyzed for allergens. Allergens and their family proteins were deleted based on reviewed protein information obtained from UniProt, and 17,372 non-allergenic proteins were prepared. Furthermore, the thymic medulla provides a unique microenvironment in which virtually all the self-antigens are presented so that autoreactive T cells are eliminated from the T cell repertoire before emigrating to the periphery, thus establishing central T cell tolerance. mTECs play a pivotal role in self-antigen presentation process by virtue of their promiscuous expression of tissue-restricted proteins. Therefore, such a protein expressed in mTECs basically cannot induce allergic reactions. The published gene and protein expressions were integrated to create 1,534 non-allergenic proteins. Fig. S1 shows the distributions of several physico-chemical features of allergenic and non-allergenic proteins, where, for each protein, these feature values are computed by the average for amino acids in it, with the feature values for each amino acid being presented in Data S3. There is no significant difference in the distributions of any physicochemical features between them, which suggests that it is impossible to predict allergenicity by simply using those features.

Fig. 1(**C**) shows examples of amino-acid sequences and patterns (subsequences). We call contiguous amino-acid subsequences of various lengths *patterns*. The goal of this study is to find the patterns highly associated with allergic reactions as ASPs. The challenges stem from the fact that there exists an extremely large number of candidate patterns to be considered. For example, if we consider patterns up to length 50, because there are 20 amino-acids, there are 20 + 20^2^ + … + 20^50^ = 10^65^ possible patterns. In fact, we have only to consider patterns contained in the dataset, but even so, the total number of patterns in the PoC dataset is as large as 3, 783, 825, 994. Such an extremely large number of candidate patterns causes not only computational but also statistical difficulties. The main contribution in this paper is to develop a method for overcoming these computational and statistical difficulties. The number of patterns contained in the PoC dataset by their lengths is shown in Fig. S2.

### Allergen-Specific Patterns (ASPs)

In this study, we call the patterns that satisfy all the following three conditions as ASPs. The definition of an ASP is that

1. the pattern is observed statistically significantly more frequently in allergenic proteins than in non-allergenic proteins,
2. the pattern is observed in none of the non-allergenic proteins, and
3. the pattern is not specific to particular biological category.

The first condition is needed to guarantee the statistical reliability of the identified ASPs. The second condition means that if an ASP is the cause of the allergic reaction, it should not be observed in any non-allergenic proteins. The third condition suggests that patterns specific to a particular biological category should not be mistakenly identified as ASPs. In this study, the third condition is verified by checking whether one of the following conditions is met:

3a) the pattern is observed in at least one paired category, or

3b) the pattern is observed in multiple positive-only categories.

Because a paired category contains both allergenic and non-allergenic proteins, a pattern observed more frequently in allergenic proteins can be considered allergen-specific, rather than specific to the biological category (condition 3a). On the other hand, because it is not possible to distinguish whether the pattern contained in a single positive-only category is specific to the biological category or the allergic reaction, we only regard the patterns as ASPs if they are commonly contained in multiple positive-only categories (condition 3b). Fig. 2(**A**) shows the definition of ASPs and Fig. 2(**B**) shows examples of ASPs and non-ASPs.

**Fig. 2.**
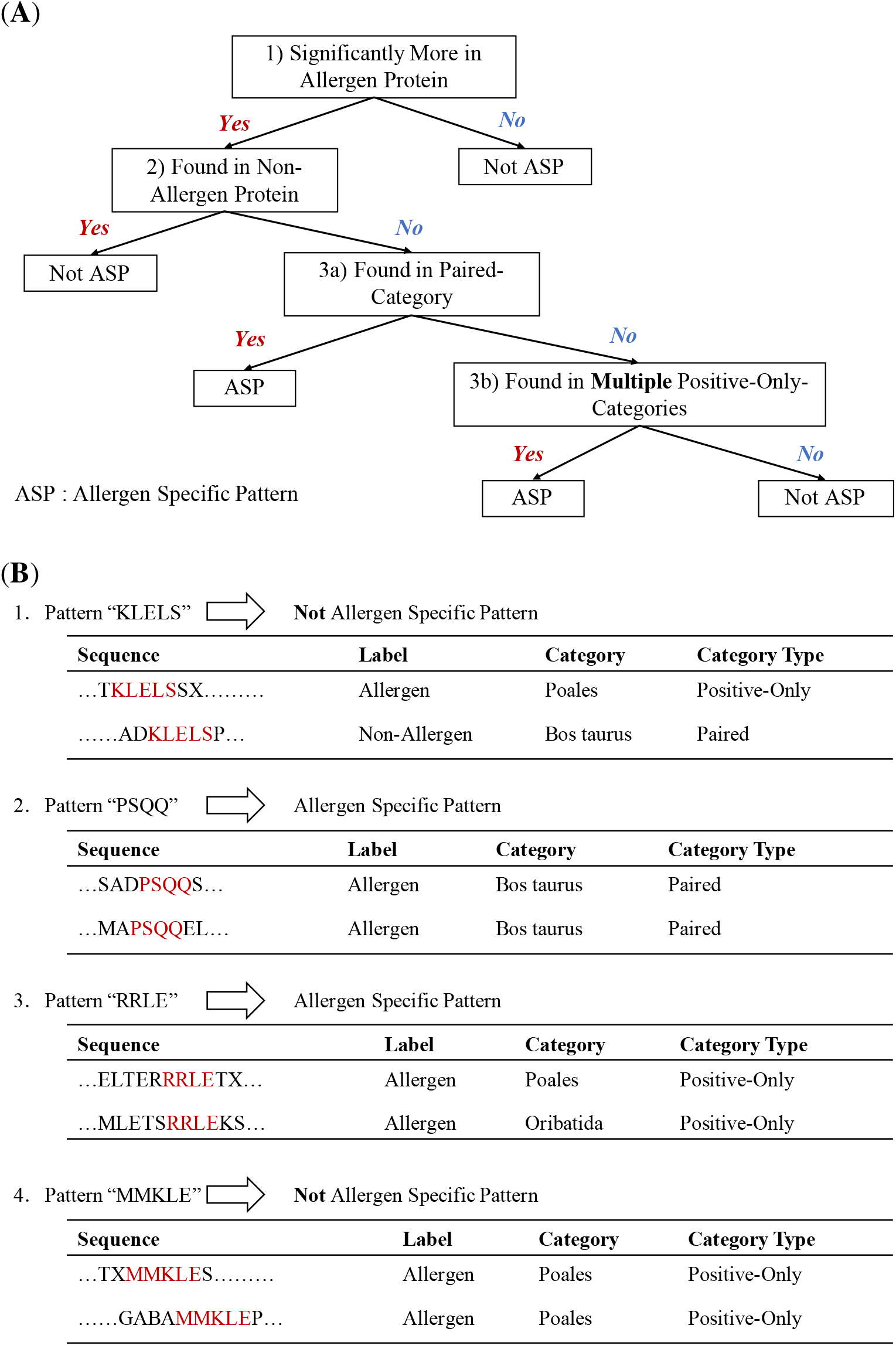
Definition and examples of allergen-specific patterns (ASPs). (**A**) Decision diagram to define ASPs. (**B**) Examples of ASPs and non-ASPs. These examples show the circumstances under which a pattern that satisfies the condition 1 is defined as an ASP. (Example 1) Pattern KLELS is not an ASP because it does not satisfy condition 2. (Example 2) Pattern PSQQ is an ASP because it satisfies conditions 2 and 3a. (Example 3) Pattern RRLE is an ASP because it satisfies conditions 2 and 3b. (Example 4) Pattern MMKLE satisfies condition 2, but is not an ASP because it is found only in a single positive-only category, i.e., satisfies neither condition 3a nor 3b.

### Statistically Significant Pattern Mining Method

Because the number of candidate patterns is huge, it is necessary to introduce a computational trick to reduce the computational cost. Efficient algorithms for handling large number of patterns have been studied extensively in the data mining community. In particular, methods for efficiently handling sequential patterns is called sequence mining (*17–24*), and our method is built on one of sequence mining methods called *prefix-span* (*20*). The large number of candidate patterns raises not only a computational challenge but also a statistical one. Over the past decade, several methods have been proposed for evaluating the statistical significance of discovered patterns in several data mining tasks (*33, 45–49*). In this paper, we evaluate the statistical reliability of ASPs by adapting the techniques developed in these studies to our problem setup.

As mentioned above, because the number of candidate patterns is huge, the computational cost will be extremely large if all the candidate patterns are explicitly handled in the method. In sequence mining, a tree structure of subsequence patterns is constructed, and efficient computation is made possible by pruning the branches of the tree structure, as illustrated in Fig. 3(**A**). In the most basic sequence mining task called frequent sequence mining, one can efficiently find patterns whose frequency (called *support*) is greater than a certain threshold. Frequent sequence mining takes advantage of the fact that the frequency of the parent pattern in the tree structure is no less than the frequency of the child patterns. Fig. 3(**A**) illustrates the concepts of tree structure and pruning in a frequent sequence mining task.

**Fig. 3.**
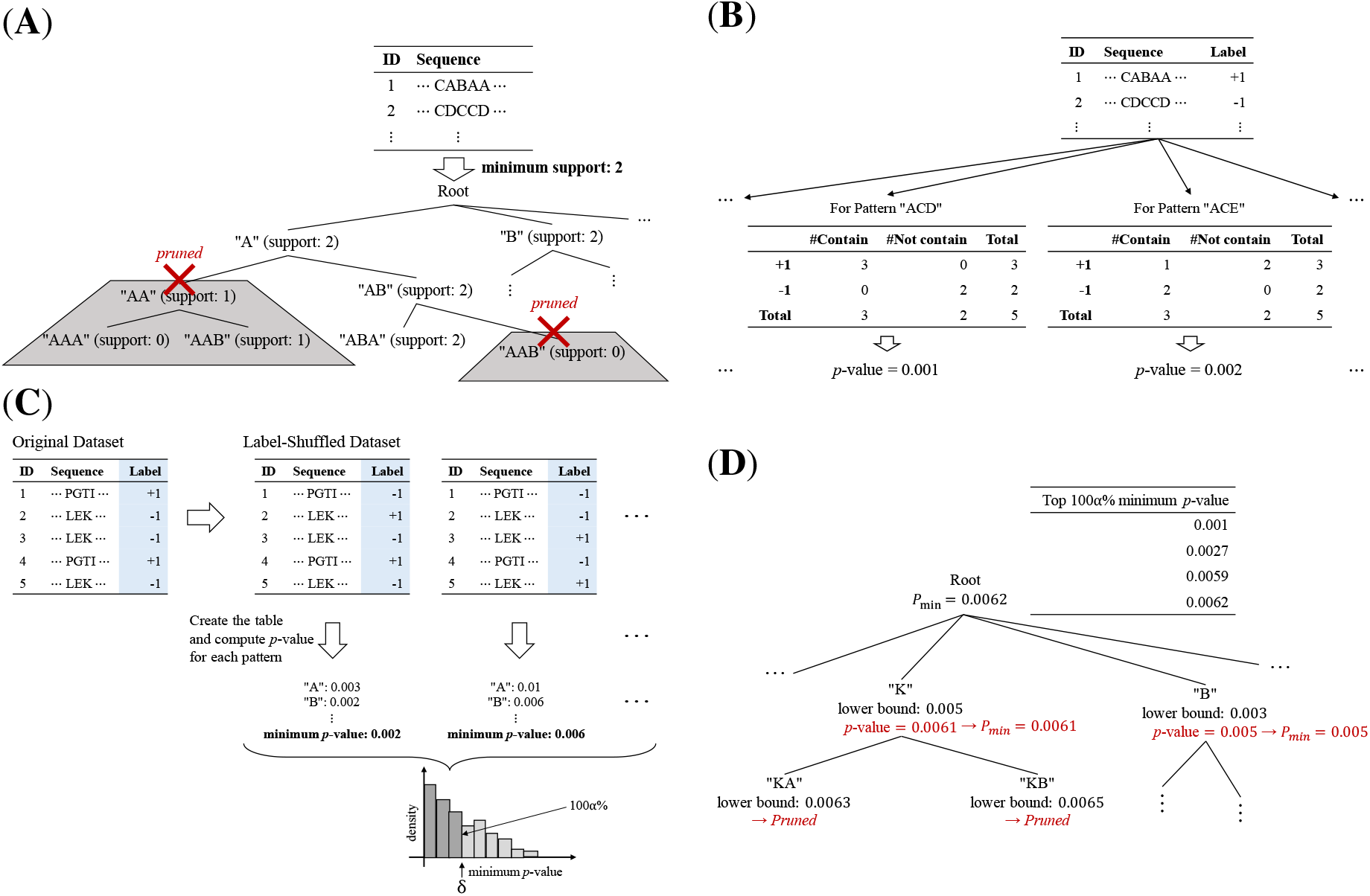
Schematic illustrations of the proposed method. (**A**) Illustration of a sequence mining problem and tree pruning. (**B**) Illustration of multiple testing problem with multiple Fisher exact test each of which corresponds to each pattern. (**C**) Family-wise error rate (FWER) controlled by the Westfall-Young (WY) method. (**D**) Illustration of a fast computation of WY method by exploiting the tree structure and its pruning.

Statistical hypothesis testing based on contingency tables can be used to statistically evaluate whether certain amino-acid subsequence is more frequently observed in allergenic proteins. We employed the Fisher Exact Test (FET) for testing the significance of a contingency table. Because we consider a large number of candidate patterns, we need to consider testing the statistical significance of a large number of contingency tables by multiple FETs. When multiple tests are conducted, a multiple testing correction is needed to adequately control the risk of false positive (FP) findings. In the context of multiple testing, one is often required to control the probability of finding one or more FPs below the significance level, a criterion known as the *family-wise error rate (FWER)*. The most basic multiple testing correction method for controlling FWER is called the Bonferroni correction, for which one needs to use the significance level divided by the number of tests as the adjusted significance level. Unfortunately, in our problem, the number of candidate patterns (i.e., the number of hypotheses) is so large that the Bonferroni correction is over-conservative. In fact, because there are more than 3.7 billion candidate patterns in the PoC dataset, only those with the (nominal) FET *p*-value smaller than 0.05*/*3, 783, 825, 994 < 1.3 × 10^−11^ can be considered statistically significant for FWER < 0.05. Fig. 3(**B**) illustrates that the statistical reliabilities of the identified ASPs can be quantified by considering multiple testing problems with multiple FETs each of which corresponds to each pattern in the dataset. Various multiple testing correction and other selection bias correction methods have been studied (*25–32, 34–37*).

In this study, we introduce a method for multiple testing correction for large number of contingency tables by effectively combining a randomized test called the *Westfall-Young (WY)* method (*50*) with a sequence mining method. In WY method, the labels (i.e., allergen/non-allergen) are randomly shuffled to create a randomized dataset which does not contain any ASPs by construction. Because all selected patterns from the shuffled dataset in WY method are interpreted as FP findings, to avoid them, the significance threshold for FWER control must be smaller than all the (nominal) FET *p*-values obtained from the shuffled datasets. By generating multiple (e.g., 10,000) shuffled datasets with different random seeds, we can estimate the distribution of minimal (nominal) FET *p*-values. The adjusted significance level is defined to be the lower 100*α*% point of the minimal (nominal) FET *p*-value distribution. Fig. 3(**C**) schematically illustrates the multiple testing correction for evaluating the statistical reliability of the selected ASPs by WY method.

Unfortunately, because the WY method requires the calculation of (nominal) FET *p*-values for all possible patterns, it cannot be used as it is. Thus, we introduce a trick to avoid computations that does not affect the lower 100*α*% of the minimal (nominal) FET *p*-value distribution. Concretely, we compute the lower bound of the (nominal) FET *p*-values of the patterns included in each branch of the tree structure shown in Fig. 3(**C**). Then, it is possible to avoid the computation for the patterns which do not affect the lower 100*α*% point of the minimum (nominal) FET *p*-value distribution by pruning the branches based on the lower bound. The main technical difference of allerStat from existing sequence mining methods is that the tree pruning strategy is designed so that branches only containing patterns not affecting the estimation of the FWER distribution in the WY method are pruned. This computational trick was first introduced in (*46*), in which the goal was to quantify the statistical significance of itemset mining tasks. In this study, we adapted the method to our sequence mining task.

### Statistical Properties of the ASPs in the PoC dataset

The ASPs identified in the PoC dataset are listed in Data S4. In total, 5,064 and 5,994 statistically significant ASPs were found when the statistical significance thresholds were set to be *α* = 0.01 and 0.05, respectively. Figs. 4(**A**), 4(**B**), and 4(**C**) show the distributions of the lengths, the adjusted *p*-values, and the supports of the identified ASPs, respectively. Fig. 4(**D**) shows the distributions of the number of ASPs in each allergenic protein. Fig. 4(**E**) shows the distribution of how many different biological categories each ASP is included in.

**Fig. 4.**
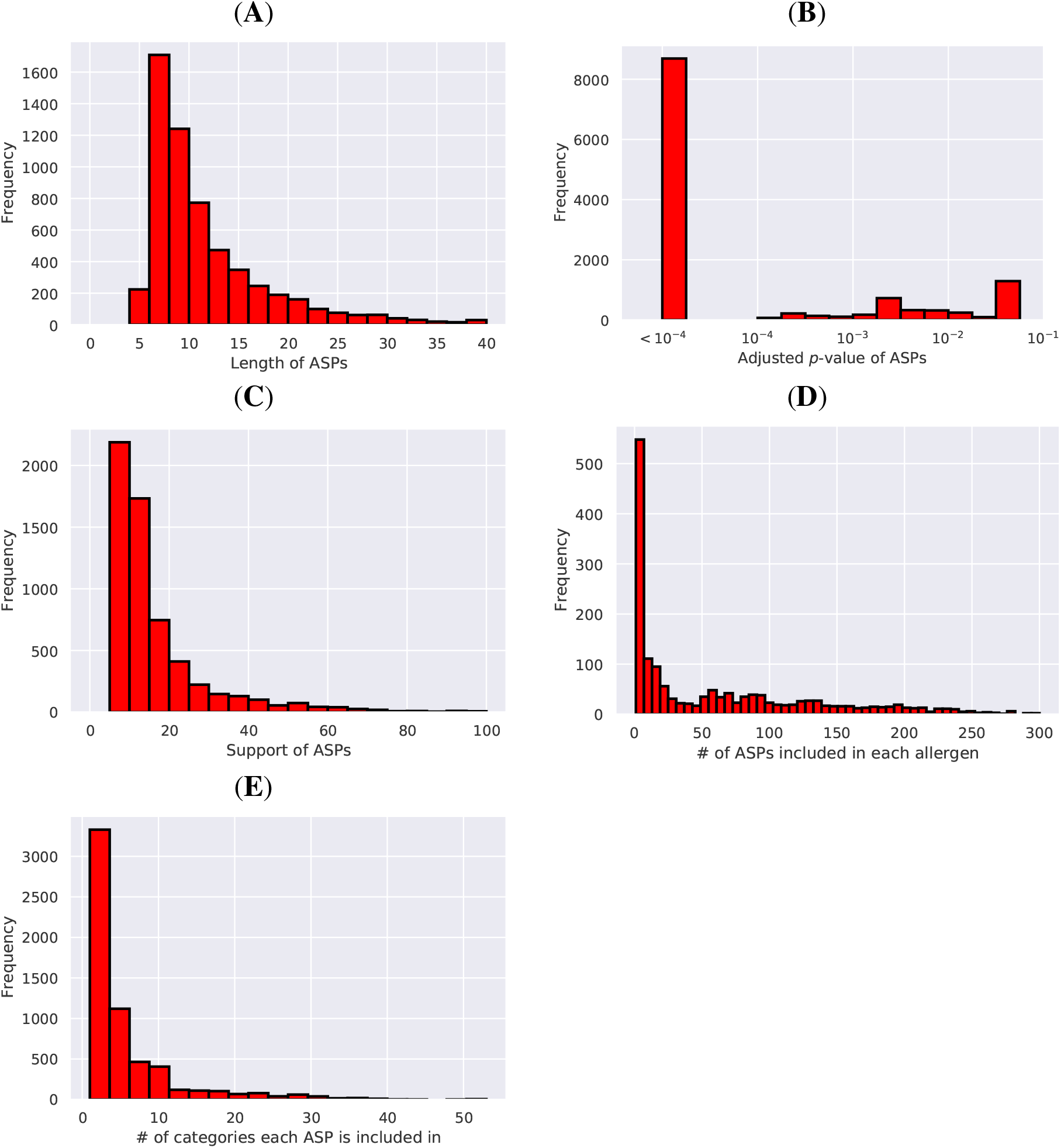
Statistical properties of allergen-specific patterns (ASPs) in the proof-of-concept (PoC) dataset. (**A-C**) Distributions of the lengths, the adjusted *p*-values, and the supports of the identified ASPs, respectively. (**D**) Distributions of the number of ASPs in each allergenic proteins. (**E**) Distribution of how many different biological categories each ASP is included in.

### Biological Analysis of ASPs

The 5,994 ASPs detected by allerStat with *α* = 0.05 were reprocessed to remove overlapped sequences and were concatenated to 1,072 patterns (ConcASPs; Data S5) to make it easier for further analysis (see the Methods section and Supplementary Text 1). Among the ConcASPs, 687 patterns had a length of 15 amino acids or more. Therefore, we next examined HLA type II binding activities. A number of recent studies suggest that HLA plays a significant role in atopic diseases and food allergies (*51–53*). Protein antigens in foods are internalized by antigen presenting cells (APCs) such as dendric cells and B cells. These antigens are processed into short peptides (12–25 amino-acid long) that can bind to HLA-II. Antigen-specific T cells recognize peptide-HLA-II complex, leading to allergic reactions. The binding to HLA-II molecules, such as HLA-DR, is considered as the first step to antigen-specific IgE antibody production. The association of the ConcASP and typical 15 HLA-DR alleles such as HLA-DRB1 was investigated using NetMHCIIpan 4.0 (*38*). A total of 82% of the 687 ConcASPs with ≥ 15 amino-acid long carried a HLA-DRB1 binding activity we examined (Fig. 5(**A**)). Of the 687 ConcASPs, 40.8% for HLA-DRB1*13.01 to 59.1% for HLA-DRB1*04.01 showed HLA-DRB1 binding activity for each major allele. Consensus motifs of core sequences to each HLA-DRB1 allele are shown in Fig. 5(**B**) as Web logo. The full result of the existences of binding for the 687 ConcASPs and 15 HLA-DR alleles is presented in Data S6.

**Fig. 5.**
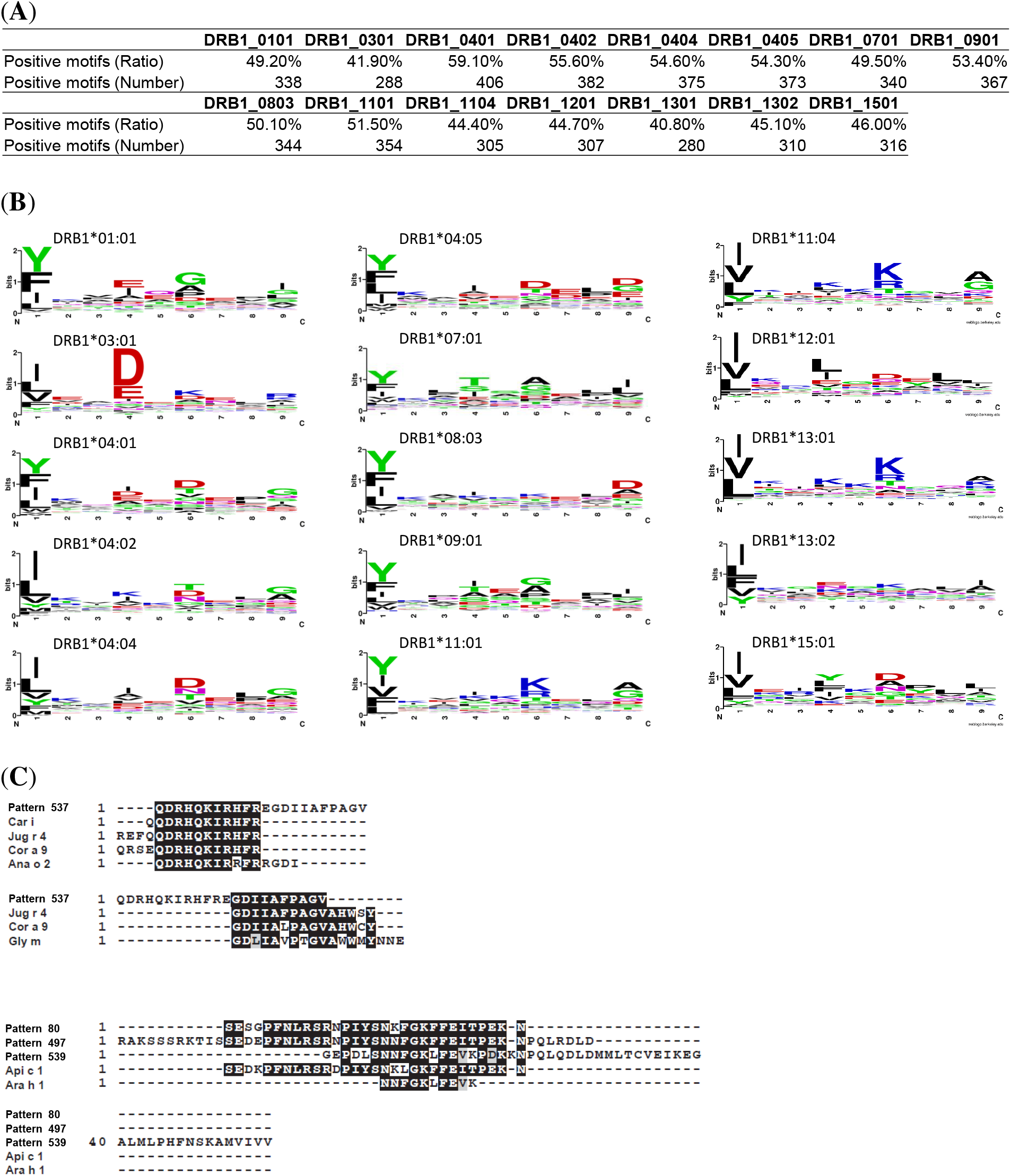
Biological characteristics of the sequence patterns specific to allergenic proteins. Protein sequences that have amino acids with a length of 15 or more were examined to predict human leukocyte antigen (HLA)-DRB1 binding activities. (**A**) Percentage of HLA binding motifs contained in the extracted sequence patterns. (**B**) Consensus motifs of core sequences to each HLA-DRB1 allele are shown as sequence logos. (**C**) ConcASP patterns contain validated B-cell epitope sequence. BLAST search was performed to find similarity with B-cell epitope.

Furthermore, we examined the relationship between identified ASPs and known IgE epitopes. The web server program called allergen database for food safety (ADFS) provides analytical tools for searching the similarity with these validated B-cell epitopes for which site binds to the IgE from patients with food allergy. We performed a homology search with online alignment search program called Protein Basic Local Alignment Search Tool (BlastP). As a result, 24.3% of ConcASP were highly homologous to the sequences of B-cell epitopes (Data S7). For instance, pattern 537 had a high degree of sequence homology with seven epitopes identified from tree nut and bean allergens, suggesting an allergen cross reactivity (Fig. 5(**C**)). Remarkably, patterns 80, 497, and 539 shared homologies with epitope sequences identified in two completely different species: bee and peanut. Some more specific examples are presented in Fig. S3. Additionally, 225 out of 1072 ConcASP (21.0%) were perfectly matched with the known food allergen epitopes.

### Predicting Allergic Reaction using ASPs

We considered the problem of predicting allergic reactions of unknown proteins using the ASPs identified by allerStat. Allergic reaction prediction from amino-acid sequences is needed for safety assessments of biotechnology-based synthetic foods such as genome editing.

Cross-validation (CV) is often employed in the evaluation of predictive modeling. CV evaluates predictive performance by removing some instances from the dataset, training an ML model with the remaining instances, and applying the trained model to the removed instances for performance evaluation. It is important to note that CV is based on the assumption that each instance is independently identically distributed (i.i.d.). Because the proteins used in this study are strongly associated within the same biological category, we used what we call *leave-category-out CV (LCO-CV)*. In LCO-CV, all proteins in each category were removed, a machine learning model is trained with the proteins in the remaining categories, and the prediction performance is investigated by applying the trained model to the proteins in the removed category. We considered LCO-CV for each paired category. Fig. 6(**A**) is a schematic illustration of LCO-CV.

**Fig. 6.**
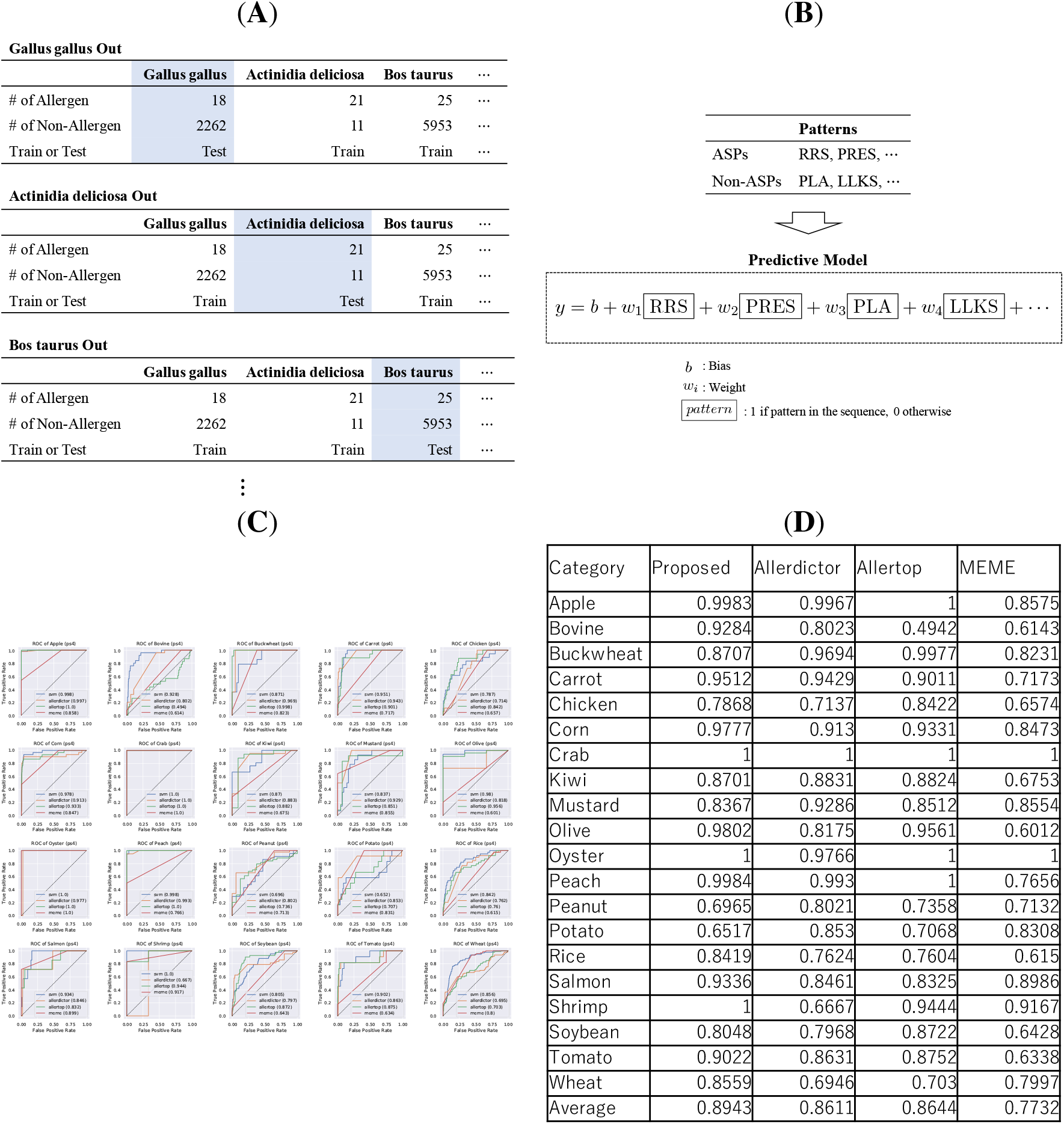
Illustrations and results of prediction analysis. (**A**) Schematic illustration of leave-category-out cross validation (LCO-CV). (**B**) Schematic illustration of prediction analysis by sparse support vector machine (SVM). (**C**) Receiver operating characteristic (ROC) curves for 11 paired categories in LCO-CV. (**D**) Area-under-the-curve (AUC) scores for 11 paired categories in LCO-CV.

To predict the presence or absence of an allergic reaction, it is desirable to use not only the ASPs but also the patterns that are more frequently observed in non-allergens. Therefore, we also considered non-Allergen-Specific Patterns (non-ASPs) which were similarly identified as ASPs. Non-ASPs differ from ASPs in that non-ASPs should be significantly more frequently appeared in non-allergenic proteins, but may also be appeared in a small number of allergenic proteins. Therefore, without the condition corresponding to the second condition 2) in the definition of ASP, a non-ASP is defined as

1′) the pattern is observed statistically significantly more frequently in non-allergenic proteins than in allergenic proteins, and

3′) the pattern is not specific to particular biological category.

When predicting an allergic reaction of a new protein, it is desirable to interpret why the protein is or is not determined to have allergic reaction. Some ML models, such as deep neural networks, are called black-box models in that are too complex to interpret how the predictions are made. Because the interpretation of the prediction process is important in the current problem, we adopted a linear prediction model in which a feature is defined by the existence of each ASP or non-ASP in the amino-acid sequence. In the training set, each row corresponds to a protein and each column corresponds to each ASP or non-ASP, where the ASPs and non-ASPs in the training set are patterns that were identified without using the proteins in the removed category by LCO-CV. Fig. 6(**B**) illustrates the concept of a predictive model based on the ASPs and the non-ASPs selected by SVM. See the Methods section for the detail of prediction model development by SVM.

Figs. 6(**C-D**) show the results of the predictive analysis based on the LCO-CV for the PoC dataset. To evaluate the effectiveness of the ASPs identified by allerStat, we compared the prediction performances with the methods proposed in previous studies. There are several previous studies on ML-based allergen prediction from amino-acid sequences (*10–16, 42–44*). An ML-based allergen prediction model consists of two steps: Step 1) feature extraction from amino-acid sequences, and Step 2) model training based on the extracted features. Because the goal of the current study is to evaluate the effectiveness of the ASPs identified by allerStat, we only compared the feature extraction method at Step 1 with those of previous studies, and used the same ML model (sparse SVM) at Step 2.

We compared allerStat with *Alledictor* (*43*), *MEME* (*42*), and *Allertop* (*44*). The features extracted in Alledictor are amino-acid subsequences as in allerStat. The main drawback of Alledictor is that it could only consider patterns of fixed length (only the patterns of the length 6 were considered in (*43*) as well as in the current experiment). MEME is a method for extracting frequent amino-acid subsequences based on a probabilistic model called the *hidden Markov model (HMM)*. The main drawback of the feature extraction method by MEME is that it is an unsupervised method and therefore cannot extract patterns observed at different frequencies in allergenic and non-allergenic proteins. Allertop differs from allerStat, Alledictor and MEME in that the features extracted by Allertop are constructed from the physico-chemical features of the entire amino-acid sequence. The main drawback of the feature extraction method by Allertop is that it is difficult to interpret which parts of the amino-acid subsequence are associated with the allergic reaction. Note that, unlike allerStat, the statistical reliabilities of the patterns extracted in these three previous methods cannot be properly evaluated.

Fig. 6(**C**) shows the receiver-operating-characteristic (ROC) curves of the prediction results based on LCO-CV. When interpreting the results, it is important to note that the sample size of each category varies greatly, and the numbers of allergenic and non-allergenic proteins are highly unbalanced. Fig. 6(**D**) shows the area-under-curve (AUC) scores of various prediction models. It can be observed that the prediction performances of allerStat are consistently better than the three existing methods. The information on the prediction model trained with the entire PoC dataset is provided in Data S8.

## Discussions

The problem of finding ASPs based on a database of allergenic/non-allergenic proteins can be regarded as a typical feature selection problem for a binary classification problem. However, it is important to note that the proteins in the database are not i.i.d., which is a common assumption taken in conventional classification problem. In particular, proteins belonging to the same category (apple, bovine, etc.) have amino-acid sequences that are specific to that category, and if the search is not conducted carefully, category-specific amino-acid sequences will be identified instead of allergen-specific ones. To address this problem, we divided the proteins in the database into categories and used different criteria for ASPs depending on whether each category was paired, positive-only, or negative-only. Furthermore, in the allergen prediction task, care must be taken not to make predictions based on category-specific amino-acid sequences by introducing LCO-CVs in which each category is excluded from the training data. In the protein allergen study, it is important to carefully take into account for the “non i.i.d.-ness” of proteins.

Our results show that 82% of the sequence patterns specific to allergenic proteins obtained by allerStat carried the binding activity to any of the 15 major HLA-DRB1 alleles. Almost 50% of the patterns had the HLA-DRB1 binding affinity for either of the 15 HLA-DRB1 alleles. This indicates that allerStat can extract biologically significant patterns. Sequence logo analysis indicates that the core sequences had characteristic motifs at the position of 1, 4, 6, and 9 in 9 core sequences, especially DRB1*01.01, *04.01, *04.02, *07.01, *11.01 (Fig. 5(**B**)). This suggests that our extracted patterns include many core sequences important for major HLA-DRB1 binding. We did not evaluate HLA binding activities of the concASPs having a length less than 15 amino acids. The remaining 18% of the patterns that do not have any HLA-DRB1 binding activity need to be investigated to clarify their importance, because they may include biologically significant sequences. On the other hand, non-allergenic patterns have sequences with three or six amino-acid long, which do not have any consensus motif.

Among the identified ASPs, 1,083 allergen-specific IgE epitopes on 247 food allergens have been experimentally elucidated. Among the obtained 1,072 ConcASPs from allerStat, 225 of which (21%) included any of the entire length of 125 epitope sequences. On the other hand, only 19 epitope sequences were found in 17,372 non-allergen sequence (0.1%). Although three epitopes including Pen c 3 epitope (ISSK and YGVA) and Fag e 1 epitope (QQPGQ) had overlapping in between, all 19 epitopes length is in three to seven amino acids considering to have low specificity. Moreover, 24.3% of ConcASP showed high homology with known food allergen epitopes. Interestingly, there are many patterns homologous with epitopes among different species (Fig. 5(**C**)). These results indicated that allerStat could properly extracted B-cell epitope sequences characteristic to the allergen from our dataset, even though epitope information was not held in the training data. The remaining 75% patterns are thought to contain T-cell epitopes and derived unexpected sequences, such as those related to immunological tolerance and antibody production, in addition to unknown T- and B-cell epitopes.

## Methods

### Allergen and non-allergen data collection

Allergen data were retrieved from full-length amino-acid sequence list in the COMprehensive Protein Allergen Resourse (COMPARE) 2020 data (https://comparedatabase.org/). The COMPARE database, a collaborative effort of the HESI Protein Allergenicity Technical Committee, is a curated database comprising allergen-sequence-associated peer-reviewed publications. Non-allergen data were collected by reviewing protein sequence expressed in twenty foods including Wheat (Triticum aestivum), Bovine (Bos taurus), Chicken (Gallus gallus), Soybean (Glycine max), Crab (Scylla serrata), Shrimp(Penaeus monodon), Peanut (Arachis hypogaea), Buckwheat (Fagopyrum esculentum), Salmon (Salmo salar), Kiwi (Actinidia deliciosa), Mustard (Sinapis alba), Olive (Olea europaea), Carrot (Daucus carota), Apple (Malus domestica), Tomato (Solanum lycopersicum), Peach (Prunus persica), Potato (Solanum tuberosum), Corn (Zea mays) Rice (Oryza sativa), and Oyster (Crassostrea gigas), obtained from UniProt (https://www.uniprot.org/). Since these agricultural and marine products were well analyzed as allergic foods, non-allergen data was created by deleting known allergens contained in each and the proteins belongs to the same family, from whole protein data. Known allergen were listed from ADFS and COMPARE databases. mTEC data set were generated by integrating gene and protein expression profiles (*54,55*). Data reduction of gene expression in SGLT1+ mature mTECs (GEO# GSE49625) were selected with cutoff level of *t* = 5.

### HLA-II binding activity of motifs specific to allergenic proteins

Among 1,072 concatenated allergen-specific patterns, 687 of which had a length of 15 amino acids or more. HLA-DRB1 binding activity was predicted by NetMHCIIpan server ver.4 (https://services.healthtech.dtu.dk/service.php?NetMHCIIpan-4.0), which requires amino-acid sequences with 15 amino acids or longer. The HLA-DRB1 binding activity of the allergen-specific sequences were examined at 15 amino-acid windows from the N-terminus to the C-terminus. Allergen-specific patterns having strong and weak binding activities were analyzed using sequence logo to clarify specific motifs to each HLA-DRB1 allele using Web Logo (https://weblogo.berkeley.edu/logo.cgi).

### Similarity between the identified ASPs and known IgE-epitopes

Sequence similarity was searched by running BlastP with following parameters: E value = 2,000 and cutoff of 10^−4^ (https://allergen.nihs.go.jp/ADFS/). The sequences exactly matched targets were extracted from BlastP results (E value: 20,000). As a preprocessing for this purpose, we concatenated ASPs in the following criteria. For two sequences *x* and *y*, let *x* ⋄ *y* indicates the concatenated sequence. We define *overlapped concatenation* between two sequences *x* and *y* as *x*′ ⋄ *w* ⋄ *y*′ if there exists *x*′, *y*′, *w* ∈ 𝒮 such that *x* = *x*′ ⋄ *w* and *y* = *w* ⋄ *y*′. Note that the overlapped concatenation is just the ordinary concatenation *x* ⋄ *y* when we take *w* as an empty sequence. The overlapped concatenation of three or more sequences is defined as the repetition of an overlapped concatenation for two sequences. Then, we define a sequence *x* as a *concatenated ASP* (ConcASP) if (i) there exists an allergen sequence *y* in the dataset such that *x* is included in *y* as a subsequence (denoted by *x* ⊑ *y*), and (ii) *x* is represented as an overlapped concatenation of any number of ASPs. Finally, after retrieving all ConcASPs, we extracted only the maximal ones, that is, we excluded those that are included in other concatenated ASPs. In order to find completely matching IgE epitopes to concatenated ASPs, we simply examined the inclusion, that is, checked whether *x* ⊑ *y* for each *x* in IgE epitope sequences and each *y* in ConcASPs. Here, an IgE epitope may consist of multiple sequences, and it may contain non-sequence motifs. So we extracted only sequence motifs from them. As a result, 1,104 sequences were retrieved from 247 epitopes. Among the 1,072 concatenated ASPs, 225 completely matched at least one epitope sequence. We present the algorithm for overlapped concatenation as Algorithm 1 in Supplementary Text 1.

### Fisher Exact Test (FET)

Let *D, D*^+^, and *D*^−^ represent the sets of all proteins, allergenic proteins, and non-allergenic proteins, respectively. Furthermore, the sizes of *D, D*^+^, and *D*^−^ are respectively written as *n, n*^+^, and *n*^−^. To quantify whether there is a difference in the frequencies of patterns between allergenic and non-allergenic proteins, we consider the following contingency table,

| | Containing<br>pattern $q$ | Not containing<br>pattern $q$ | Sum |
| --- | --- | --- | --- |
| Allergen | $D^+[q]$ | $n_+ - D^+[q]$ | $n_+$ |
| Non-allergen | $D^-[q]$ | $n_- - D^-[q]$ | $n_-$ |
| Sum | $D[q]$ | $n - D[q]$ | $n$ |

where *D*g[*q*], *D*^+^[*q*], and *D*^−^[*q*] denote the number of proteins in *D, D*^+^, and *D*^−^ which contains pattern *q*. To test the statistical significance, Fisher exact test (FET) is used. The *p*-value of FET is computed by considering all possible realizations of the contingency table under the condition that *n*^+^, *n*^−^ and *D*[*q*] are fixed. Specifically, let

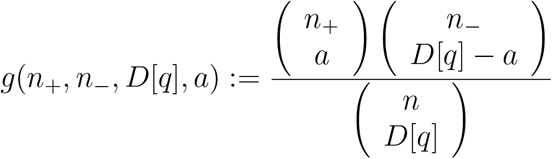

be the probability of finding a pattern *q* in *a* of *n*_+_ allergenic proteins conditional on *n*^+^, *n*^−^ and *D*[*q*]. The two-tailed FET *p*-value is then calculated as

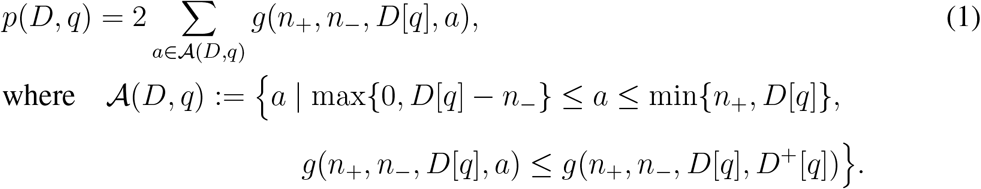

### Multiple testing correction by Westfall-Young (WY) method

When we conduct only one hypothesis test, the type-I error, i.e., the probability of FP finding, can be controlled at a significance level *α* ∈ (0, 1) (typically *α* = 0.05) by rejecting the null hypothesis when the *p*-value is less than *α*. However, when we conduct many hypothesis tests simultaneously, we often need to control the FWER: the probability of having one or more FP findings. To control the FWER at the significance level of *α*, we need to reject the null hypothesis more conservatively, i.e., a null hypothesis is rejected when the *p*-value is less than the adjusted significance level *δ* ∈ (0, 1) where *δ* is usually much smaller than *α*. Methods for finding the adjusted significance level *δ* such that the FWER can be controlled to *α* is discussed in the context of multiple comparisons. The most commonly used multiple comparison method is Bonferroni correction in which the adjusted significance level *δ* is set to be *α/*{# of hypotheses}, where {# of hypotheses} is the number of hypotheses to be tested simultaneously. Bonferroni correction is known to be overly conservative, especially when the number of hypotheses is large. In allerStat, we employ a random permutation test called the Westfall-Young (WY) method (*50*) as a multiple test correction method. In the WY method, multiple randomized datasets are constructed by permuting the labels (allergenic or non-alleregenic in our problem setup) at random. For each randomized dataset, the minimum *p*-value is computed. Then, the adjusted significance level *δ* is set to be the ⌈(*α* + 1)*/M* ⌉^th^ smallest value among *M* smallest *p*-values where *M* is the number of randomized datasets (typically *M* = 10000). Fig. 3(**C**) illustrates the notion of WY method.

### Sequence mining and efficient computation of the WY method

Unfortunately, since aller-Stat needs to handle an extremely large number of patterns, it is impossible to calculate the minimum *p*-value for each of the *M* randomized datasets for the WY method. In allerStat, we consider a tree structure among patterns as shown in Fig. 3 for efficiently computing the minimum *p*-value. Specifically, we consider a node corresponding to a pattern *q* and the set of its descendant sequence patterns 𝒮 (*q*). Then, our basic idea is to compute a lower bound of the FET *p*-values of the descendant sequence patterns in the form of

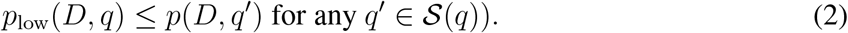

If we can find such a lower bound, we can reduce the computational cost because we do not need to calculate the *p*-values of all descendant nodes when we encounter a node whose lower bound *p*_low_(*D, q*) is greater than the current minimum *p*-value. Here, we employed a lower bound that satisfies the property in (2), written as

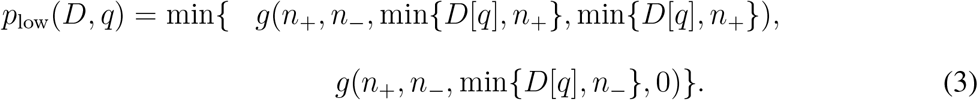

The lower bound in (3) was proposed in FastWY method in (*45, 46*), in which the authors studied itemset mining tasks to find statistically significant combinations of multiple genetic factors. In allerStat, we adapted the technique in the FastWY method for sequence mining setting. Therefore, the proof that the bound (3) satisfies the property in (2) can be simply done as described in (*45, 46*).

We present the algorithm for AllerStat as Algorithm 2 in Supplementary Text 2.

### Prediction Model for Allergic Reaction using ASPs and non-ASPs

In order to predicting allergic reactions by proteins using ASPs and non-ASPs, we employed the support vector machine (SVM) with features being defined by ASPs and non-ASPs. We follow the formulation of SVM employed by *LinearSVC* class in *scikit-learn* (https://scikit-learn.org/stable/modules/generated/sklearn.svm.LinearSVC.html), a python implementation that we used. For an overview of SVM, see, for example, Chapter 7 of (*56*).

SVM assumes that we are given *n* instances, where each instance consists of *d* input variables **x**_*i*_ ∈ **R**^*d*^ and binary output variable *y*_*i*_ ∈ {−1, 1} (1 ≤ *i* ≤ *n*). Here, **R** denotes the set of all real numbers, and **R**^*d*^ the set of all *d*-dimensional real vectors. Then, given a test instance whose *d*-dimensional input variables are **x**^*′*^ while its output variable is unknown, suppose that its prediction result is given by the linear model

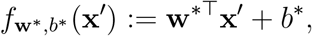

where **w**^*^ ∈ **R**^*d*^ and *b*^*^ ∈ **R** are to be learned by SVM. If *f*_**w**\*,*b*\*_ (**x**^*′*^) is larger than (resp. smaller than) zero, the prediction function conjectures that the output variable for **x**^*′*^ is 1 (resp. −1). From the training dataset 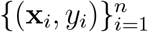 defined above, SVM learns optimal **w**^*^ and *b*^*^ by

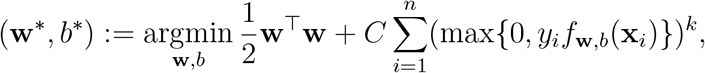

where *C* is a hyperparameter to control overfitting or underfitting, while *k* ∈ {1, 2} is an option of learning. For *C*, in this computational experiment we chose the best one in AUC by LFO-CV from log_10_ *C* ∈ {−27*/*9, −23*/*9, −19*/*9, …, 9*/*9} (10 cases). Note that, since this LFO-CV is conducted for the training set composed by an LFO-CV, this LFO-CV for the selection of *C* is conducted for a remained set of biological categories. For *k, k* = 1 is the original formulation of SVM, while *k* = 2 is a modification for computational efficiency (*57*). In these computational experiments we used *k* = 2.

Then we show how to apply ASPs by allerStat and protein sequences to the formulation of SVM above. Let 𝒬_*j*_ ∈ 𝒮 (1 ≤ *j* ≤ *d*) be an ASP or a non-ASP, *z*_*i*_ ∈ 𝒮 (1 ≤ *i* ≤ *n*) be a training sequences, and *y*_*i*_ ∈ {−1, 1} (1 ≤ *i* ≤ *n*) be their allergenicity (1 if allergenic or −1 otherwise). Then we define **x**_*i*_ ∈ **R**^*d*^ as the indicator variables of including patterns in *z*_*i*_, that is, *x*_*ij*_ = 1 if 𝒬_*j*_ ⊑ *z*_*i*_ or *x*_*ij*_ = 0 otherwise. As a result, we can employ SVM to learn a allerginicity prediction function according to ASPs and non-ASPs. Here, we can see that the *j*^th^ element in **w**^*^ is the importantness of the corresponding pattern: if it is a large positive number, the pattern is expected to contribute the allergenicity. Note that the training sequences are defined by LFO-CV stated in the Results section.

Finally we show how the existing methods are compared to allerStat stated in the Results section. For Allerdictor and MEME, we employed the same process as that for allerStat, except for the retrievals of patterns. For Allerdictor, we used all length-six patterns rather than ASPs and non-ASPs. For MEME, since it is an unsupervised method, we ran MEME by providing only allergenic proteins to retrieve ASPs, then by providing only non-allergenic proteins to retrieve non-ASPs. For Allertop, since it provides a fixed-length vector from a protein sequence, we just used it for the SVM training and the prediction.

## Supporting information

Zipped Supplementary Information

## Funding

This work was supported in part by the following funds:

The Ministry of Health, Labor and Welfare grant H30-Syokuhin-Ippan-002 (NT, KK, IT)

The Ministry of Health, Labor and Welfare grant 21KA1002 (KK)

MEXT KAKENHI grant 20H00601 (IT)

JST Moonshot R&D grant JPMJMS2033-05 (IT)

NEDO grants JPNP18002, JPNP20006 (IT)

RIKEN Center for Advanced Intelligence Project (IT)

## Author contributions

Conceptualization: NT, RA, KK, IT

Methodology: KG, TY, HH, TS, IT

Resources: NT, RA, KK

Software: KG, TY, HH, TS, IT

Visualization: KG, NT, TY, HH, KK

Writing — original draft: KG, NT, HH, KK, IT

## Data and materials availability

All the data are provided as Supplementary Data. The software is available at https://github.com/takeuchi-lab/allerStat.

