## Supplementary material for "AllerStat: Finding Statistically Significant Allergen-Specific Patterns in Protein Sequences by Machine Learning": Zipped Supplementary Information: supp.pdf

**This PDF file includes:**

Supplementary Texts S1 and S2

Figs. S1 to S3

**Other Supplementary Materials for this manuscript include the following:**

Data S1 to S8

### Supplementary Text S1: Overlapped concatenation of ASPs

The algorithm of finding concatenated ASPs is presented as Algorithm 1.

### Supplementary Text S2: Algorithm of AllerStat

In this supplementary text, we formally describe the proposed AllerStat method. In allerStat, we consider a tree structure shown in Fig. 3 to solve the computational and statistical challenges caused by considering an extremely large number of patterns (amino-acid subsequences). The tree representation of patterns enables us to take advantage of a branch-and-bound technique by finding a condition such that, if a condition holds for a node in the tree, its descendant nodes cannot be ASPs. As described previously, there are three conditions to be met in order for a pattern to be an ASP. Our basic strategy is to exploit the tree structure to efficiently search for patterns that satisfy condition 1, and then to check if they satisfy conditions 2 and 3. The method for finding patterns that satisfy condition 1 is built on Fast Westfall-Young (FastWY) method (*46*), which was developed for finding statistically significant combinations of multiple genetic factors by effectively combining an itemset mining method with a random permutation testing method. Here, we extended the basic techniques of FastWY method to a sequence mining task for identifying amino-acid sequences which satisfy condition 1, i.e., which appears significantly more frequently in allergenic proteins than in non-allergenic proteins. We present the algorithm of AllerStat in Algorithm 2.

**Algorithm 1** Calculating concatenated ASPs

**Require:** $A \subset \mathcal{S}$, $F \subset \mathcal{S}$.
**Ensure:** $C \subset \mathcal{S}$.
  $\{A$: all allergen sequences, $F$: all ASPs, $C$: all concatenated ASPs$\}$
**for** $i \in \{1, 2, \ldots, |A|\}$ **do**
  Initialize $B_k \leftarrow 0$ $(1 \in k \in |A_i|)$.
  $\{$Binary variables to check appearances of ASPs in allergen sequences$\}$
  **for** $j \in \{1, 2, \ldots, |F|\}$ **do**
    **if** $F_j \sqsubseteq A_i$ **then**
      Let $m$ be the position of $F_j$ found in $A_i$.
      $\{m := |w| + 1$ if $w \diamond F_j \diamond w' = A_i$.$\}$
      **for** $m$: all positions of $F_j$ found in $A_i$ **do**
        $B_{ik} \leftarrow 1$ for $k = m, \ldots, m + |F_i| - 1$.
      **end for**
    **end if**
  **end for**
  **for** $1 \leq b < e \leq |A_i|$ **do**
    **if** $B_b = B_{b+1} = \ldots = B_e = 1$ and $B_{b-1} = B_{e+1} = 0$ **then**
    $\{$For convenience, define $B_0 = B_{|A_i|+1} = 0\}$
      $C \leftarrow C \cup \{(A_i)_{b..e}\}$
    **end if**
  **end for**
**end for**
Exclude any sequence in $C$ such that it is included by another sequence in $C$.
**return** $C$.

**Algorithm 2** Algorithm of allerStat

**Require:** $\alpha \in (0,1)$, $\tilde{D}^{+}, \tilde{D}^{-} \subseteq \mathcal{S}$, $M > 0$.
 {$\alpha$: significance level in FWER, $\tilde{D}^{+}$: positive (allergen) sequences,}
 {$\tilde{D}^{-}$: negative (non-allergen) sequences, $M$: number of trials of WY}
**Ensure:** $\delta \in (0,1)$, $Q^{+} \subseteq \mathcal{S} \times (0,1)$, $Q^{-} \subseteq \mathcal{S} \times (0,1)$.
 {$\delta$: significance level for single hypothesis testing,}
 {$Q^{+}$: ASPs and their $p$-values, $Q^{-}$: non-ASPs and their $p$-values}

 {Run FastWY algorithm}
Prepare $(L_1, L_2, \ldots, L_M)$. {The smallest $p$-value among multiple tests in the $M^{\mathrm{th}}$ trial}
**for** $m \in \{1, 2, \ldots, M\}$ **do**
  Let $\ddot{D}^{+}$ and $\ddot{D}^{-}$ be the shuffled dataset of $\tilde{D}^{+}$ and $\tilde{D}^{-}$, on condition that
   $|\ddot{D}^{+}| = |\tilde{D}^{+}|$ and $|\ddot{D}^{-}| = |\tilde{D}^{-}|$.
  $L_m \leftarrow$ MinimumPValue($\ddot{D}^{+}$, $\ddot{D}^{-}$). {Described in Algorithm 3.}
**end for**
Let $\delta$ be the next smaller value than the $\lceil (\alpha + 1)/M \rceil$th smallest value among
 values in $L_m$ ($m = 1, 2, \ldots, M$).
$Q \leftarrow$ SignificantPatterns($\delta$, $\tilde{D}^{+}$, $\tilde{D}^{-}$).
 {Find patterns whose $p$-values without shuffling are less than $\delta$.}
 {Described in Algorithm 3.}

 {Select patterns that are ASPs or non-ASPs in accordance with Results section}
$Q^{+} \leftarrow \{\}$; $Q^{-} \leftarrow \{\}$. {Set of ASPs and non-ASPs, respectively}
**for** $(q, p) \in Q$ **do** {$q$: pattern, $p$: its $p$-value}
  **if** sequences $\{x \mid x \in \tilde{D}, q \sqsubseteq x\}$ are from only one positive-only category **then**
   **continue** {$q$ is category-specific rather than allergen-specific}
  **end if**
  **if** $\tilde{D}^{-}[q]/|\tilde{D}^{-}| > \tilde{D}^{+}[q]/|\tilde{D}^{+}|$ **then**
  {Since we conduct two-sided FET, $Q$ can contain both ASPs and non-ASPs.}
  {This condition extracts only non-ASPs (more frequent in non-allergens).}
   Add $(q, p)$ to $Q^{-}$.
  **else if** $\tilde{D}^{-}[q] = 0$ **then**
   Add $(q, p)$ to $Q^{+}$. {ASPs must not be observed in non-allergens}
  **end if**
**end for**
**return** $Q^{+}$.

**Algorithm 3** Enumeration of patterns and the pruning conditions of AllerStat

**function** MINIMUMPVALUE($\ddot{D}^{+}$, $\ddot{D}^{-}$)
{Computes the pattern having the minimum $p$-value, and returns the $p$-value.}
  $Q \leftarrow$ ENUMPATTERNS(**null**, $\ddot{D}^{+}$, $\ddot{D}^{-}$).
  **return** the $p$-value in $Q$.
**end function**

**function** SIGNIFICANTPATTERNS($\delta$, $\tilde{D}^{+}$, $\tilde{D}^{-}$)
{Computes patterns whose $p$-value is $\delta$ or less.}
  **return** ENUMPATTERNS($\delta$, $\tilde{D}^{+}$, $\tilde{D}^{-}$).
**end function**

**function** ENUMPATTERNS($\delta$, $D^{+}$, $D^{-}$) {$\delta =$ **null** when called from MINIMUMPVALUE}
  $Q \leftarrow \{\}$.
  $p_{min} \leftarrow 1$. {Currently smallest $p$-value}
  $q_{min} \leftarrow$ “”. {Pattern that provides $p_{min}$}
  $T \leftarrow \{c \mid c \in \mathcal{S}, |c| = 1\}$. {Set of all patterns of length 1}
  **while** $T \neq \emptyset$ **do**
    Remove an element in $T$ and store to $q$.
    Calculate $p_{\text{low}}(D, q)$ in (3).
    **if** $p_{\text{low}}(D, q) \leq (\delta \mid\mid p_{min})$ **then** {$a$ || $b$: “$a$ if it is not null, $b$ otherwise”}
      **continue** {$q$ is pruned}
    **end if**
    Calculate $p(D, q)$ in (1).
    **if** $\delta \neq$ **null** and $p(D, q) \leq \delta$ **then**
      Add $(q, p(D, q))$ to $Q$.
    **end if**
    **if** $p(D, q) < p_{min}$ **then**
      $p_{min} \leftarrow p(D, q)$;  $q_{min} \leftarrow q$.
    **end if**
    $T \leftarrow T \cup \{q \diamond c \mid c \in \mathcal{S}, |c| = 1\}$.
  **end while**
  **if** $\delta =$ **null then** $Q \leftarrow (q_{min}, p_{min})$.
  **return** $Q$.
**end function**

**Fig. S1.** Distributions of physicochemical features of allergenic and non-allergenic proteins in the PoC dataset, where, for each protein, these feature values are computed by the average for amino acids in it, with the feature values for each amino acid being presented in Data S3.

**Fig. S2.** The number of patterns contained in the PoC dataset by their lengths (up to length 50). In total there are 3,783,825,994 patterns.

**Fig. S3.** Biological characteristics of the sequence patterns specific to allergenic proteins.

**Fig. S1 (1)**

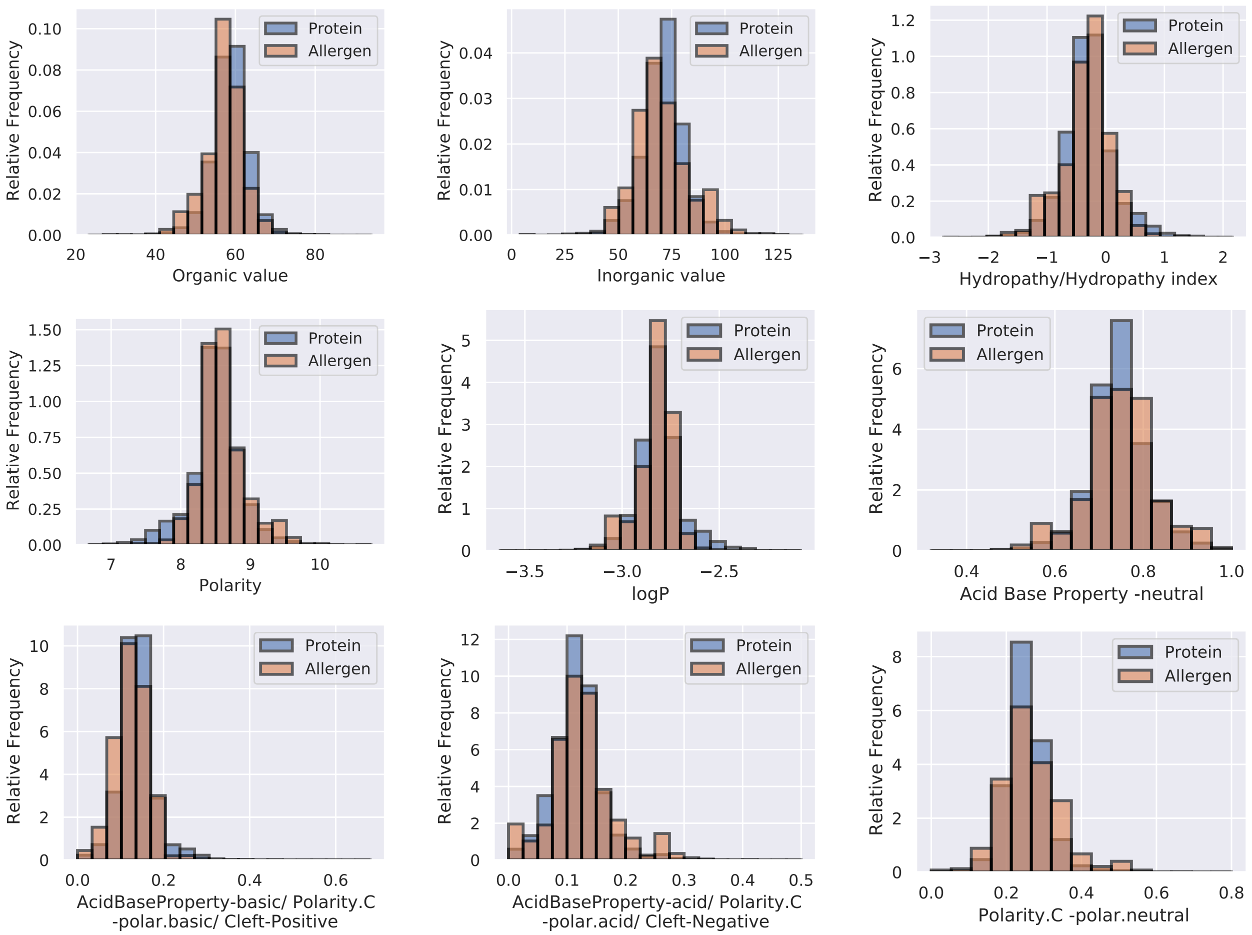

Relative Frequency
Protein
Allergen
Organic value
Inorganic value
Hydropathy/Hydropathy index
Polarity
logP
Acid Base Property -neutral
AcidBaseProperty-basic/ Polarity.C -polar.basic/ Cleft-Positive
AcidBaseProperty-acid/ Polarity.C -polar.acid/ Cleft-Negative
Polarity.C -polar.neutral

**Fig. S1 (2)**

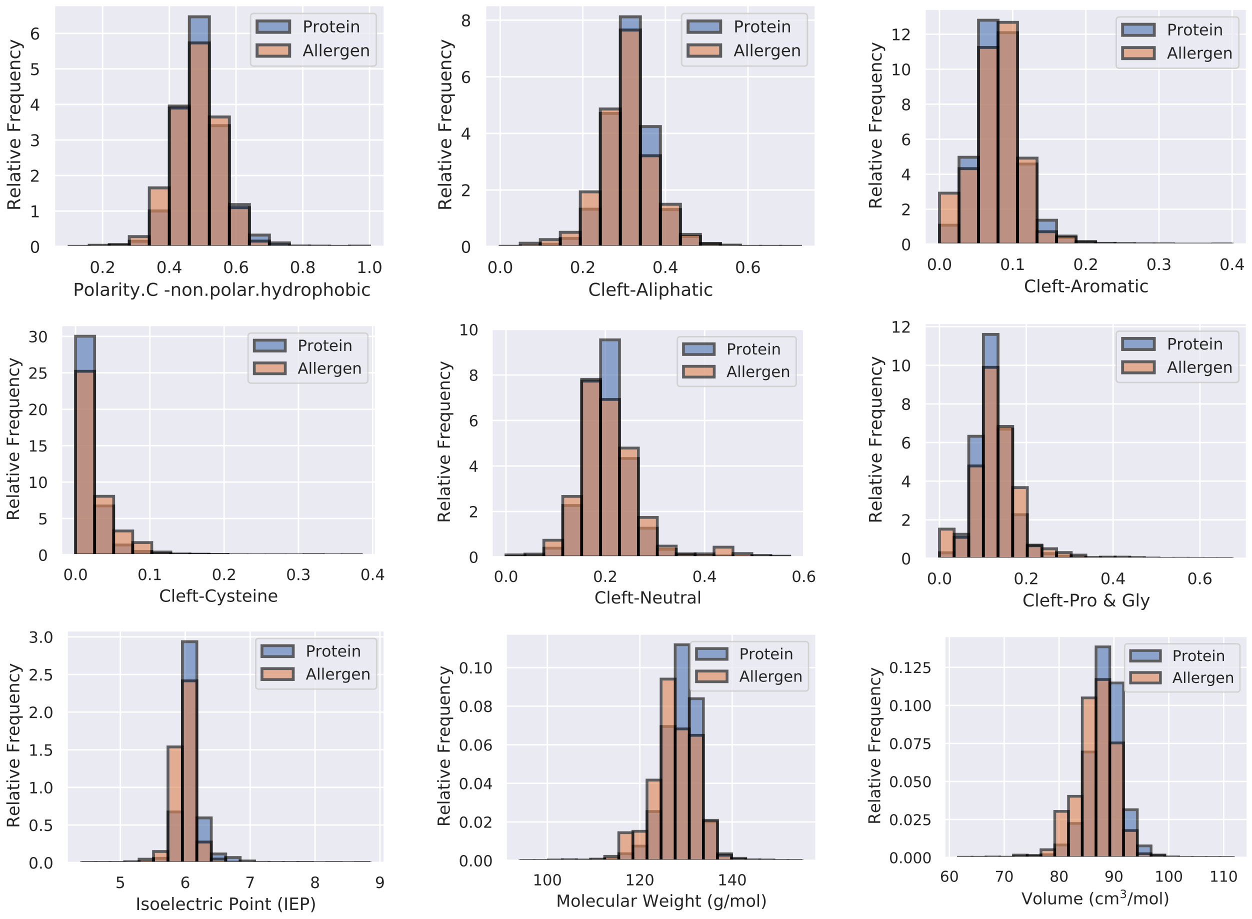
Relative Frequency
Protein
Allergen
Polarity.C -non.polar.hydrophobic
Cleft-Aliphatic
Cleft-Aromatic
Cleft-Cysteine
Cleft-Neutral
Cleft-Pro & Gly
Isoelectric Point (IEP)
Molecular Weight (g/mol)
Volume (cm³/mol)

**Fig. S2**

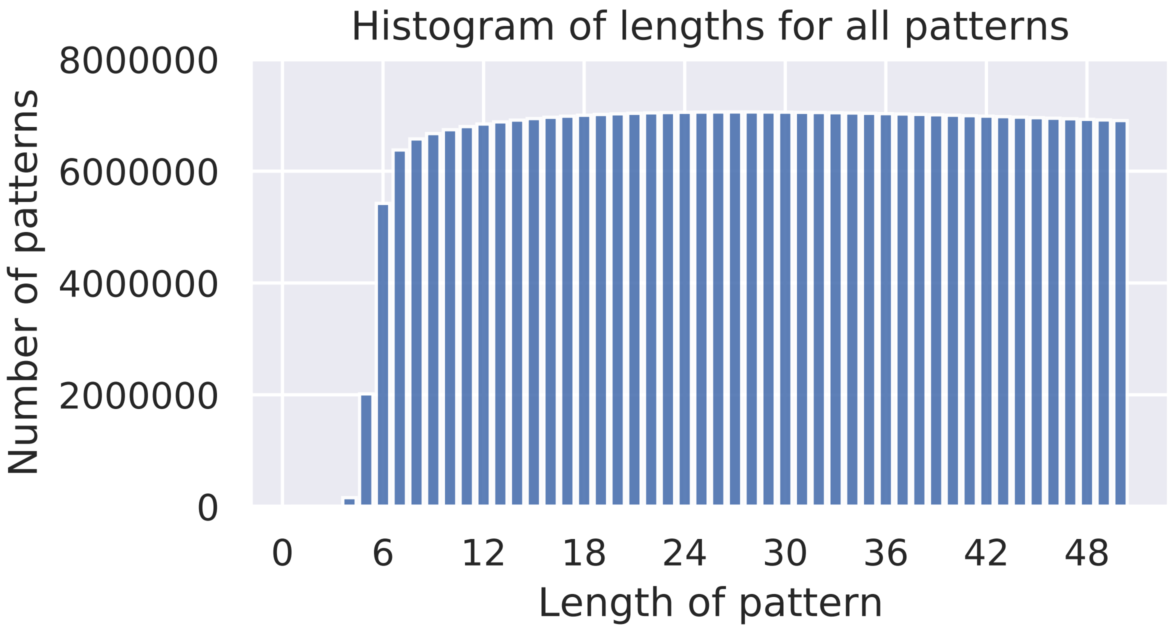

Histogram of lengths for all patterns
8000000
6000000
4000000
2000000
0
Number of patterns
0
6
12
18
24
30
36
42
48
Length of pattern

**Fig. S3**

#### Bovine and Goat

```
Pattern 538   1 QQQPPFSQQQPILPQQPPFSQQQQLVLPQQSPFSQQQQ--------
Bos t         1 QSEEQQQTEDELQDKIHPFAQTQSLVYPFPGP--------------
Car h         1 ---------------IHPFAQAQSLVYPFTGPIPNSLPQNILPLTQ
```

#### Yorkshire-fog and Crayfish

```
Pattern 614   1 -------------TACGNVPIFKDGKGCGSCYE----
Hol l 1       1 GYKDVDKPPFSGMTGCGNTPIFKDGRGCGSCFEIK--
Pro c         1 ----------------------FKDRKDGSCYVSYKV
```

#### Chicken and Cedar

```
Pattern 688   1 ALGDTLEKICNEIKIVATPDGGC---------
Gal d 1       1 ----------------KRHDGGCRKE------
Jun a 3       1 ------------------VDGGCNSACNVFKT
```

#### Melon and Sunflower and Crayfish

```
Pattern 718   1 VKKTGQALIFGIYDEPVTPGQCNMIVERLGDYLVEQGM
Cuc m 2       1 ------------YDEPLTPGQCNMIVE-----------
Pro c         1 ---------RIIYGGSVTPGNCKE--------------
Hel a 2       1 ------------YDEPVAPG------------------
```

#### Soybean and Bovine

```
Pattern 856   1 NQLDQFPRRFYLAGNQEQEFLRYQQQ----
Gly m         1 --------RFYLAGNQEQEFLKY-------
Bos t         1 ----------PQRDMPIQAFLLYQEPVLGP
```

#### Cladosporium and Penicillum and Pea

```
Pattern 885   1 ---------------------LSGTSMASPHIAGLL---------------------
Cla h 9       1 IFAPGQDILSAWIGSTTATNTISGTSMATPHIVGLSVYLMGLENLSGPAAVTARIKE
Pen ch 18     1 -------ILSTWVGSDHATNTISGTSMA-----------------------------
Pis s         1 --------AAHEVLSWSFHSELSGTSSSKQ---------------------------
```

**Data S1.** Entire proof-of-concept (PoC) dataset containing 2,248 allergenic proteins and 18,906 non-allergenic proteins.

**Data S2.** Category information in the PoC dataset.

**Data S3.** Physico-chemical features defined for amino acid residues used to calculate the result of Fig. S1.

**Data S4.** Identified allergen-specific patterns (ASPs) by allerStat (5,064 and 5,994 statistically significant ASPs for $\alpha = 0.01$ and $0.05$, respectively).

**Data S5.** All concatenated ASPs (ConcASPs) for $\alpha = 0.05$: Concatenated sequences in Data S4 with the manner in Supplementary Text S1.

**Data S6.** The table of whether bindings are found for all pairs of ConcASPs and HLA-DR alleles. Since the software NetMHCIIpan 4.0 provides two types of binding (strong and weak), the data also provides which binding is found.

**Data S7.** The list of ConcASPs that are homologous to the sequences of B-cell epitopes.

**Data S8.** Information on the prediction model trained with the entire PoC dataset.
